# Bayesian Connective Field Modeling: a Markov Chain Monte Carlo approach

**DOI:** 10.1101/2020.09.03.281162

**Authors:** Azzurra Invernizzi, Koen V. Haak, Joana C. Carvalho, Remco J. Renken, Frans W. Cornelissen

## Abstract

The majority of neurons in the human brain process signals from neurons elsewhere in the brain. Connective Field (CF) modeling is a biologically-grounded method to describe this essential aspect of the brain’s circuitry. It allows characterizing the response of a population of neurons in terms of the activity in another part of the brain. CF modeling translates the concept of the receptive field (RF) into the domain of connectivity by assessing the spatial dependency between signals in distinct cortical visual field areas. Standard CF model estimation has some intrinsic limitations in that it cannot estimate the uncertainty associated with each of its parameters. Obtaining the uncertainty will allow identification of model biases, e.g. related to an over- or under-fitting or a co-dependence of parameters, thereby improving the CF prediction. To enable this, here we present a Bayesian framework for the CF model. Using a Markov Chain Monte Carlo (MCMC) approach, we estimate the underlying posterior distribution of the CF parameters and consequently, quantify the uncertainty associated with each estimate. We applied the method and its new Bayesian features to characterize the cortical circuitry of the early human visual cortex of 12 healthy participants that were assessed using 3T fMRI. In addition, we show how the MCMC approach enables the use of effect size (beta) as a data-driven parameter to retain relevant voxels for further analysis. Finally, we demonstrate how our new method can be used to compare different CF models. Our results show that single Gaussian models are favoured over differences of Gaussians (i.e. center-surround) models, suggesting that the cortico-cortical connections of the early visual system do not possess center-surround organisation. We conclude that our new Bayesian CF framework provides a comprehensive tool to improve our fundamental understanding of the human cortical circuitry in health and disease.

**Highlights:**

□ We present and validate a Bayesian CF framework based on a MCMC approach.
□ The MCMC CF approach quantifies the model uncertainty associated with each CF parameter.
□ We show how to use effect size *beta* as a data-driven threshold to retain relevant voxels.
□ The cortical connective fields of the human early visual system are best described by a single, circular symmetric, Gaussian.

## 1. Introduction

The majority of the neurons in the human brain process and integrate signals from neurons elsewhere in the brain (Robinson, 1989). The resulting spatial and temporal interactions and integration result in cortical feedback and feedforward mechanisms that underlie key brain functions such as human perception (Calvert & Thesen, 2004; Liang et al., 2017).

Over the past decades, the extensive use of functional magnetic resonance imaging (fMRI) combined with the rapid development of biologically-grounded computational models allowed for better understanding the relation, dependency and function of cortical areas of the human visual system (Adaszewski et al., 2018; Benson & Winawer, 2018; Dumoulin & Wandell, 2008; Meindertsma et al., 2017; Park et al., 2002; Wandell & Wade, 2003; Wandell & Winawer, 2015; Zeidman et al., 2018). One possible approach to disentangle this network of cortico-cortical interactions along the human visual pathway is connective field (CF) modeling (Haak et al., 2013). CF modeling allows characterizing the cortical response properties of a population of neurons in terms of the activity in another region of the visual cortex. It translates the concept of the stimulus-referred receptive field (RF) into the domain of connectivity by assessing the spatial dependency between signals in distinct cortical visual field areas (Haak et al., 2013). This neural-referred receptive field is also known as the cortico-cortical population receptive field (cc-pRF) and has so far been mainly used to investigate the connective plasticity in different ophthalmic diseases (Ahmadi et al., 2019; Carvalho et al., 2019; Haak et al., 2016; Haak et al., 2013; Halbertsma et al., 2019). The intrinsic advantage of using the CF is the ability to assess the spatial integration properties of the visual system using both task-evoked or restingstate neural responses (Bock et al., 2015; Gravel et al., 2014). However, the standard CF model has an intrinsic limitation. Most importantly, it does not allow for model comparisons. Enabling rigorous model comparisons for the CF model, similar to what has been done for the pRF (Zeidman et al., 2018; Zuiderbaan et al., 2012), will allow us to learn something about the neurobiology of the visual cortex through alternative hypothesis testing. Second, standard CF modeling cannot quantify the variability and reliability of each CF parameter estimated. Obtaining the uncertainty will allow the identification of model biases, e.g. related to an over- or under-fitting or a co-dependence of parameters, thereby improving the CF prediction. Finally, the model parameters obtained using standard CF modelling procedures do not lend themselves well for statistical inference at the level of single individuals, which can be important in the context of case studies, e.g. with a neurodegenerative ocular disorder, such as glaucoma.

To enable this, we here present a Bayesian inference framework for the CF modelling. Using a Markov Chain Monte Carlo (MCMC) approach (Robert & Casella, 2011), it estimates the underlying posterior distribution of the CF position and CF size, thereby quantifying the model uncertainty associated with each parameter. In addition, based on a linear spatiotemporal model of the fMRI response, we also estimate the CF effect size, which we will refer to as *beta*, adhering to the nomenclature used in Zeidman et al. (Zeidman et al., 2018). We show how this parameter can be used as a data-driven threshold to select relevant voxels. Finally, we use our new approach to perform a model comparison which demonstrates that, unlike pRFs, CFs do not appear to possess a spatial center-surround organisation.

## 2. Methods

### 2.1 Participants

Twelve healthy female participants (mean age 22 years, s.d. = 1.8 years) with normal or corrected-to-normal vision and without a history of neurological disease were included. These data were included, and used as normative dataset in previous work (Halbertsma et al., 2019). The ethics board of the University Medical Center Groningen (UMCG) approved the study protocol. All participants provided written informed consent. The study followed the tenets of the Declaration of Helsinki.

### 2.2 Stimuli presentation and description

The visual stimuli were displayed on a MR compatible screen located at the head-end of the MRI scanner with a viewing distance of 118 cm. The participant viewed the complete screen through a mirror placed at 11 cm from the eyes supported by the 32-channel SENSE head coil. Screen size was 36 x 23 degrees of visual angle and the distance from the participant’s eyes to the screen was approximately 75 cm. Stimuli were generated and displayed using the Psychtoolbox (https://github.com/Psychtoolbox-3/Psychtoolbox-3/) and VISTADISP toolbox (VISTA Lab, Stanford University), both MatLab based (Brainard, 1997; Pelli, 1997). The stimulus consisted of drifting bar apertures (of 10.2 deg radius) with a high contrast checkerboard texture on a grey (mean luminance) background. A sequence of eight different bar apertures with four different bar orientations (horizontal, vertical and diagonal orientations), two opposite motion directions and four periods of mean-luminance presentations completed the stimulus presentation that lasted 192 second. To maintain stable fixation, participants were instructed to focus on a small colored dot present in the center of the screen and press a button as soon as the dot changed color. The complete visual field mapping paradigm was presented to the participant six times, during six separate scans.

### 2.3 Data acquisition

MRI and fMRI data were obtained using a 3T Philips Intera MRI scanner (Philips, the Netherlands), with a 32-channel head coil. For each participant, a high-resolution T1-weighted three-dimensional structural scan was acquired (TR = 9.00ms, TE = 3.5ms, flip-angle = 8, acquisition matrix = 251*251*170mm, field of view = 256×170×232, voxel size = 1×1×1mm). Six VFM functional T2*-weighted, 2D echo planar images were obtained (voxel resolution of 2.5×2.5×2.5, field of view = 190×190×50 mm, TR = 1500ms, TE = 30ms). Each VFM scan lasted for 192s with a total of 136 volumes. A short T1-weighted anatomical scan with the same field of view chosen for the functional scans were acquired and used for obtaining a better co-registration between functional and anatomical volume.

### 2.4 Standard data analysis

Preprocessing and standard (pRF and CF) analysis of fMRI data were done using ITKGray (http://www.itk.org) and the mrVista (VISTASOFT) toolbox for MatLab (http://www.white.stanford.edu). The Bayesian CF approach and Bayesian CF model comparison were developed and based on MatLab 2016b (The Mathworks Inc., Natick, Massachusetts).

#### 2.4.1 Preprocessing

For each participant, the structural scan was reoriented using the anterior commissure-posterior commissure line (AC-PC line) as reference. Next, grey and white matter were automatically segmented using Freesurfer, and manually adjusted using ITKGray to minimize possible segmentation errors.

All functional data were pre-processed and analysed using the mrVista toolbox. First, motion correction within and between scans was applied. Then, we performed an alignment of functional data into anatomical space and lastly, an interpolation of functional data onto the segmented anatomical grey and white matter obtained using the ITKGray/Freesurfer.

#### 2.4.2 Population receptive field mapping

Retinotopy scans were analyzed using a model-based analysis, known as the population receptive field (pRF) method (Dumoulin & Wandell, 2008), which allows localizing the visual field maps of interest. Based on the best fitting prediction obtained using a 2D Gaussian model, the hemodynamic response (HRF), and the stimulus aperture, we estimate the visual field mapping parameters (eccentricity, polar angle and pRF size). The best model fit was projected onto a smoothed 3D mesh of the cortex, on which the visual areas were functionally identified using standard techniques (Engel et al., 1997; Sereno et al., 1994; Wandell & Winawer, 2015).

Based on the phase reversal pRF maps, all the visual cortical areas (V1, V2, V3, hV4, LO1 and LO2) were located and used to define target and source regions in the CF analysis.

#### 2.4.3 Standard connective field model

CF parameters (CF position and CF size) were estimated based on a procedure that fitted the predicted time-series in the source region (e.g. V1) to one based on the observed time-series in the target regions (e.g. V2 and V3),(Haak et al., 2013). The best fitting model parameters were retained and projected on a smoothed 3D mesh and reported on in the further analysis. The CF parameters are converted from cortical units (cortical position and size) into visual field units (eccentricity and polar angle) via a weighted integration (determined by the shape of the connective field) of the analogous pRF properties of the voxels contained by the CF in the source region.

### 2.5 Bayesian connective field model

The goal Bayesian connective field modelling is to estimate the underlying posterior distribution associated with each CF parameter, which requires an extensive search in the space of the model parameters. Here, we use a Markov Chain Monte Carlo (MCMC) approach to efficiently sample the search space. This space is spanned by the following parameters: CF center defined on a mesh grid containing the voxels of the source area, CF size, and CF effect size. Throughout this paper, we will use the abbreviation MCMC CF to indicate the new approach.

The code for the MCMC CF framework will be made available via the website www.visualneuroscience.nl.

Two MCMC CF modelling options are provided: the first option (A) estimates the standard CF parameters (position and size) using the Bayesian model. In this option the effect size (*beta*, scaling the amplitude of the predictor to the amplitude of the measured signal) is estimated using ordinary least squares fit (OSL) but it is not retained. The second option (B) estimates under joint estimation the *beta* parameter together with the other MCMC CF parameters. In this last option, *beta* is retained as meaningful and worth analysing further. The next section details the complete definition of latent variables, priors and parameters used, together with their implementation in the MCMC CF model. We will follow the nomenclature of Zeidman et al.

An overview of the MCMC CF model is presented in Figure 1 while the latent 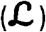 variables used are presented in table 1.

**Figure 1.**
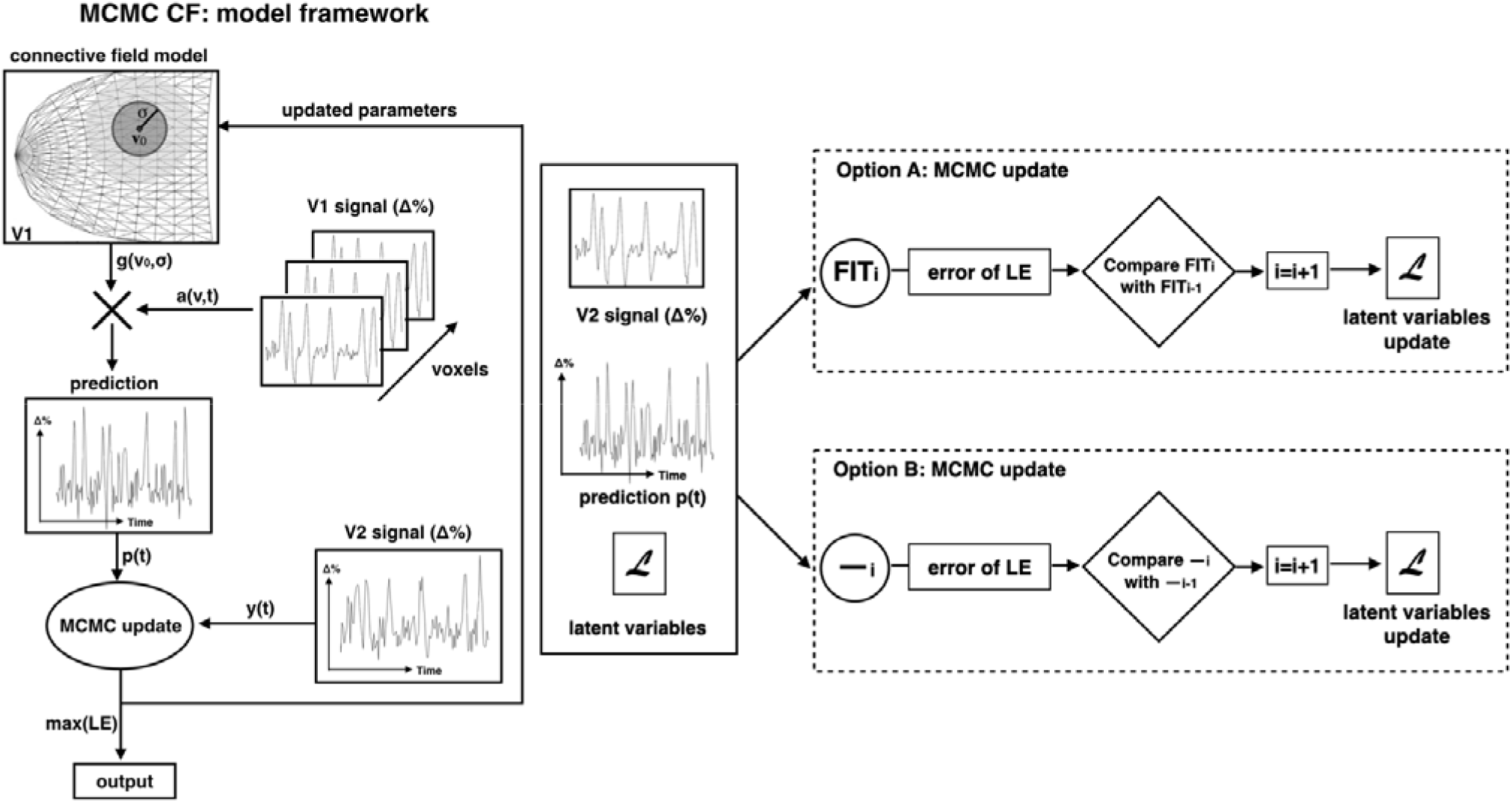
Overview of the MCMC CF framework. In this example, we estimate the V1□V2 connective field (i.e. the source region in V1 for a V2 target voxel) using the MCMC approach. Following the CF definition of Haak et al, the predicted time series is obtained by the overlap between a 2D symmetric Gaussian and the neuronal population inputs, which are the fMRI times series of all voxels in the source region (in this case V1). The predicted time series, the observed time series of the target region and the latent variables 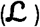 that are used to define the parameters of the connective field model. Based on the total likelihood associated with error (**LE**) calculated between and, the latent variables 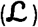 are updated in the MCMC iteration procedure (). Two different MCMC CF model options are provided: option A, in which the effect sizeis estimated using OSL (indicated as FIT in figure). While in option B, the parameter is estimated under joint estimation and retained to be further analysed - the fitting procedure used in this option is indicated as (). Figure adapted from Haak et al. (2013).

**Table 1.**
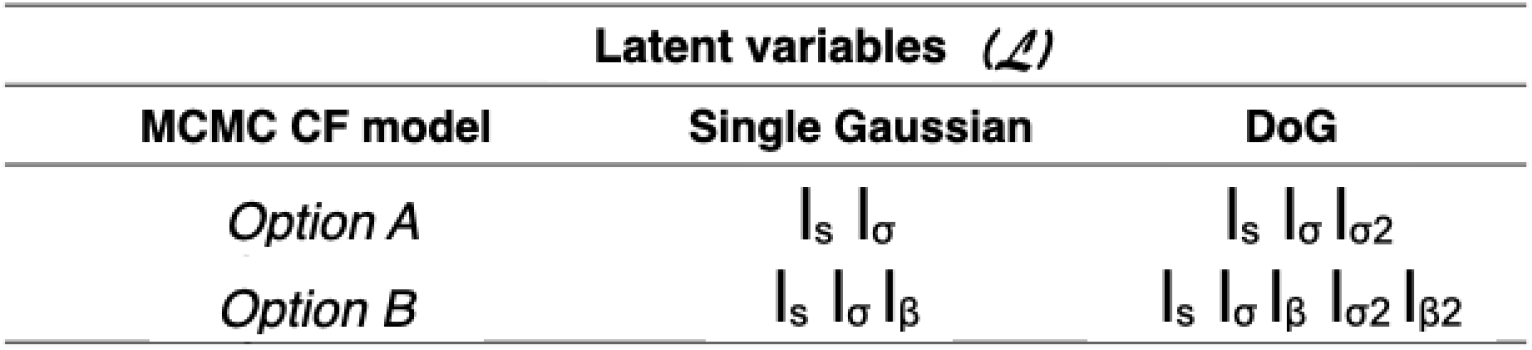
Latent variables used for the two MCMC CF options and different models implemented, respectively.

#### Latent variables and priors

Based on the CF definition used in the standard approach (Haak et al., 2013), a linear spatiotemporal model and a 2D symmetric Gaussian connective field model (2) are used to create a predicted time serie () which is fitted to the time series of a target location ().

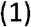

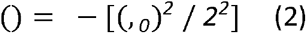

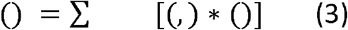

Where the predicted fMRI signal () is obtained by the overlap between the CF model() and the neuronal population inputs (,), that are defined as the BOLD time series (converted to percent signal change) for voxels () (see eq. 3). In equation 1, defines the effect size and is the error term.

The 2D symmetric Gaussian CF model of voxel (),()is defined based on the shortest three-dimensional distance (,_*0*_) between a voxel () and the proposed CF center (_0_) on a triangular mesh representation and, which defines the width of the CF.

is computed using Dijkstra’s algorithm (Dijkstra, 1959) whileis constrained to the range[_*0*_]using a latent variable(Zeidman et al., 2018). A flat prior is assumed for. Therefore, the prior for the latent variable is defined as a normal distribution (*0,1*)(see equation 4). As explained in Zeidman et al.(Zeidman et al., 2018), each latent variable is assigned to a prior distribution that represents our beliefs for that CF parameter, before the model fitting.

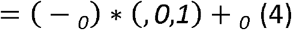

Whereis the maximum radius and _*0*_is the smallest allowed radius for the CF width - that can be an arbitrarily small non-zero number, which here were set to 10.5° and 0.01°, respectively, indicates the normal cumulative distribution function.

The MCMC is an iterative sampling approach. During each iteration the parameters for a new CF are set and the fit is compared against the current one. We will describe each parameter update in turn. A new location will be selected using the distance to the current position (). Based on the distance matrix (D), the maximum step () possible in the source region was defined as half the maximal distance from the the current position () (5). Latent variable, is randomly drawn from a normal distribution N(0,1) which results in a flat prior for the step size () between 0 and the maximum step [0] (5, 6). The updated sampling position (_*0*_) is defined as that position for which the distance to the current position is as close as possible to. If multiple locations are found, only one is drawn randomly.

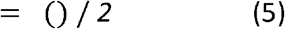

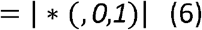

Note that for the first iteration the CF center (_*0*_) was randomly selected from the source region. Simultaneous with an updated sample location, an updated width for the CF is calculated. The is drawn from a gaussian distribution centered around the current value with a width (7).

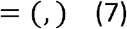

Note that in the first MCMC model (option A, Figure 1), the parameter is estimated at higher hierarchical level inside the MCMC loop using an OLS fit, i.e. theoretically, its allowed range is between –∞ and +∞. For the second MCMC CF model (option B, Figure 1), the effect size () is estimated in parallel to the other CF parameters and is retained. In this case is constrained to be positive (Zeidman et al., 2018) using the following equation:

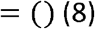

A latent variable was defined with a prior distribution (–*2,5*) and the next value was controlled by (9).

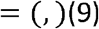

In this study, the initial values of, and were set to 1, −5 and 2, respectively.

#### MCMC CF update

At each iteration of the MCMC, the updated CF parameters (,) were estimated using the following steps. First a predicted fMRI signal() is generated from the source region using eq 2. Note that () was scaled to ensure that the total area under the gaussian, as calculated across the full source region, was equal to one. Second, the error per time point between the measured fMRI signal (()) and the predicted fMRI signal()was calculated. In MCMC CF (option A),was obtained using an ordinary least squares fit (OLS) with free parameter. While in the MCMC CF model (option B), since is estimated jointly with the other model parameters, was calculated via subtraction of the predicted signal () from the measured fMRI signal. Then, the log-likelihood associated with was estimated in the same way for both MCMC CF models (10). We assumed that follows a standard normal distribution: *N*(0,1). After estimating the mean and standard deviation of 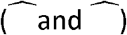, we calculated the maximum likelihood estimates (, eq. 11 and 12). Note that for MCMC CF option A only the prior for is used (11) whereas for MCMC CF option B both priors for and are included (12).

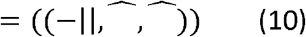

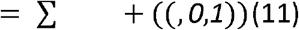

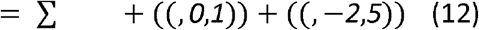

At this point, of the proposal iteration is compared to the last accepted (current) sample based on an Accepted ratio score (13).

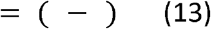

was compared to a pseudo-random acceptance score defined as a normal distribution (*0, 1*) and only if the was higher, the respective latent variables were updated. Based on the accepted, and values, a new CF was defined and a new MCMC iteration took place.

Total of 15000 iterations were run, where the first 10% of iterations were discarded for the burn-in process (Chib, 2011; Liu et al., 2016) and the posterior distributions were estimated based on the remaining part of the samples.

#### 2.5.1 Alternative kernel: Difference of Gaussians

In addition to the single gaussian model a Difference of Gaussian model (DoG) was implemented in the MCMC CF framework (Rodieck, 1965; Zuiderbaan et al., 2012). In order to do so, we integrated the two DoG model definitions proposed by Zuiderbaan et al. and by Zeidman et al. (Zeidman et al., 2018; Zuiderbaan et al., 2012). The first Gaussian is defined using the same parameters as above (i.e. *_0, 1_*, and *_1_*). The second Gaussian, which characterizes surround suppression, has the same position parameter as the first, but is defined using independent size and amplitude parameters _*2*_ and _*2*_. The size of _*2*_was enforced to be larger than_*1*_(14) while, the amplitude _2_was constrained to be negative and smaller than_*1*_(15,16). This was enforced using scaling parameters for the size and amplitudeand respectively.

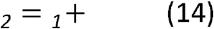

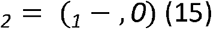

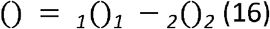

Where is constrained to be between [*0*,]using a latent variable_*2*_defined as a standard normal distribution N(0,1). is the smallest allowed radius for the second Gaussian and it was set to 0.01°. The was forced to be positive (Zeidman et al., 2018) with a latent variable _2_ with a prior distribution(–2,5).

Then, the same MCMC CF fitting procedure as described above was used. In this paper, when investigating the DoG model, we used the MCMC CF model option B in which both_*1*_and_*2*_parameters and their corresponding distributions are retained for further analysis. For the sake of completeness, the MCMC CF fitting procedure for option A is reported in supplementary material. In this study, the initial values of and were set to 5 and 10, respectively.

#### 2.5.2 CF and MCMC Model validation

For each participant, the standard CF and both MCMC CF model options were estimated using V1 as the source region that is sampled by several target regions (V2,V3,hV4, LO1 and LO2). Target and source region definitions were based on the pRF analysis described in section 2.4.2. For the purpose of model validation, variance explained was used as thresholding approach and the threshold level was chosen based on Haak et al. Only voxels for which the best-fitting CF model explained more than 15% of the time-series variance in the standard CF were included.

To quantify the level of agreement between the standard CF and each of the MCMC CF models, the parameters estimates (location and CF size) are compared using (circular) Pearson correlations at the participant and the group level for each target region.

### 2.6 Bayesian analysis and new model features

#### 2.6.1 Uncertainty

Based on a quantile analysis of the estimated posterior distribution, we computed a voxel-wise uncertainty measure (Papadopoulos & Yeung, 2001) for the MCMC CF parameters estimates as follows:

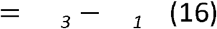

Where _*3*_and _*1*_represent the upper and lower quantiles of the posterior distribution, respectively. Furthermore, we computed the cross-correlation coefficients to quantify possible dependencies between the MCMC CF parameters (VE, CF size and *beta*; the latter only for model option B) and the associated uncertainties estimates. These correlations were computed at the participant and the group level for each target region.

#### 2.6.2 Beta thresholding approach

It is often good practice to limit further analysis of the model parameter estimates to voxels with reliable model fits. In the pRF literature this is commonly done by setting a fixed variance explained threshold (typically 15%). A more principled approach is to consider the probability that the observed model gain (or effect size) is greater than might be expected by chance alone. As such, the observed values must be compared against the null-distribution of these quantities. This null can be constructed based on theoretical distribution functions (e.g. t-distributions) and estimates of the degrees of freedom of the statistic of interest (parametric approach). However, due to both spatial and temporal dependencies, it is generally preferred to construct an empirical null by sampling the statistics of interest estimated from random data with similar spatiotemporal covariance structure (non-parametric approach).

A second consideration in this context pertains to the statistic upon which thresholding will be based. The overall model fit is intuitively quantified by the total amount of variance that is explained by the model. The threshold criterion can then be chosen such that this quantity is significantly greater than zero (e.g. the quantity falls within the 95th percentile of the null distribution). When the model involves just a single free scaling parameter (i.e. effect size), testing whether this scaling is greater than zero is often equivalent to testing the significance of the overall model fit (i.e. variance explained). This is the case in the standard pRF and CF modeling procedures as well as MCMC approach option A, where a candidate model prediction is iteratively generated and the goodness of fit is determined using a single scaling parameter (eq. 1). However under the proposed MCMC approach option B, the scaling parameter *beta* is jointly estimated with the other model parameters. Testing whether *beta* is greater than zero may therefore yield different thresholding results compared with a threshold based on residual variance. This is due to potential dependencies between the model parameters (beta and sigma). In the case of the joint estimation, parameter *beta* can be interpreted as a form of response gain (summarizing the combined effects of neuronal response gain and BOLD response). Testing the significance of *beta* for MCMC option B, therefore, provides an alternative thresholding approach that more directly addresses the question of whether or not a voxel responds to the visual stimulus in a meaningful way.

In order to test if *beta* can serve as data-driven threshold, a proxy distribution for the null hypothesis (which states that there is no correlation between the source and target region) surrogate BOLD time series were calculated for the source time-series. Based on the actual BOLD time series of each source voxel, surrogate BOLD time series were generated using the iterative amplitude adjusted Fourier transform method (iAAFT) (Räth & Monetti, 2009; Schreiber & Schmitz, 1996). A total number of 40 surrogates were computed per voxel (this number was limited due to computational feasibility). Using the iAAF method, the temporal correlations between voxels were removed while preserving their spatial relationships. Then, the MCMC CF model (B) was fitted to the surrogate BOLD time series of the source region and real time series of the target region. Using the *betas* estimated based for the surrogate time series, we derive two alternative thresholding options: 1) a *beta*-uncorrected threshold and 2) a *beta*-corrected threshold. Both *beta*-thresholds are defined using the 95th percentile of the null distribution, that can be obtained by selecting one pseudo-randomly selected surrogate or averaging all surrogates computed per voxel. For the first option, a *beta*-uncorrected threshold is obtained for each voxel in the target region. In this case, the optimal *beta*-threshold may be different for each voxel as it is empirically derived from CF fits to the surrogate BOLD time series of each individual voxel. For the second option, a *beta*-threshold is familywise error (FWE) corrected for all the voxels in the target region. In this case, one unique *beta*-threshold is obtained using the best fit of one pseudo-randomly selected surrogate BOLD time series. Figure 2 provides an overview of this *beta* thresholding approach.

**Figure 2.**
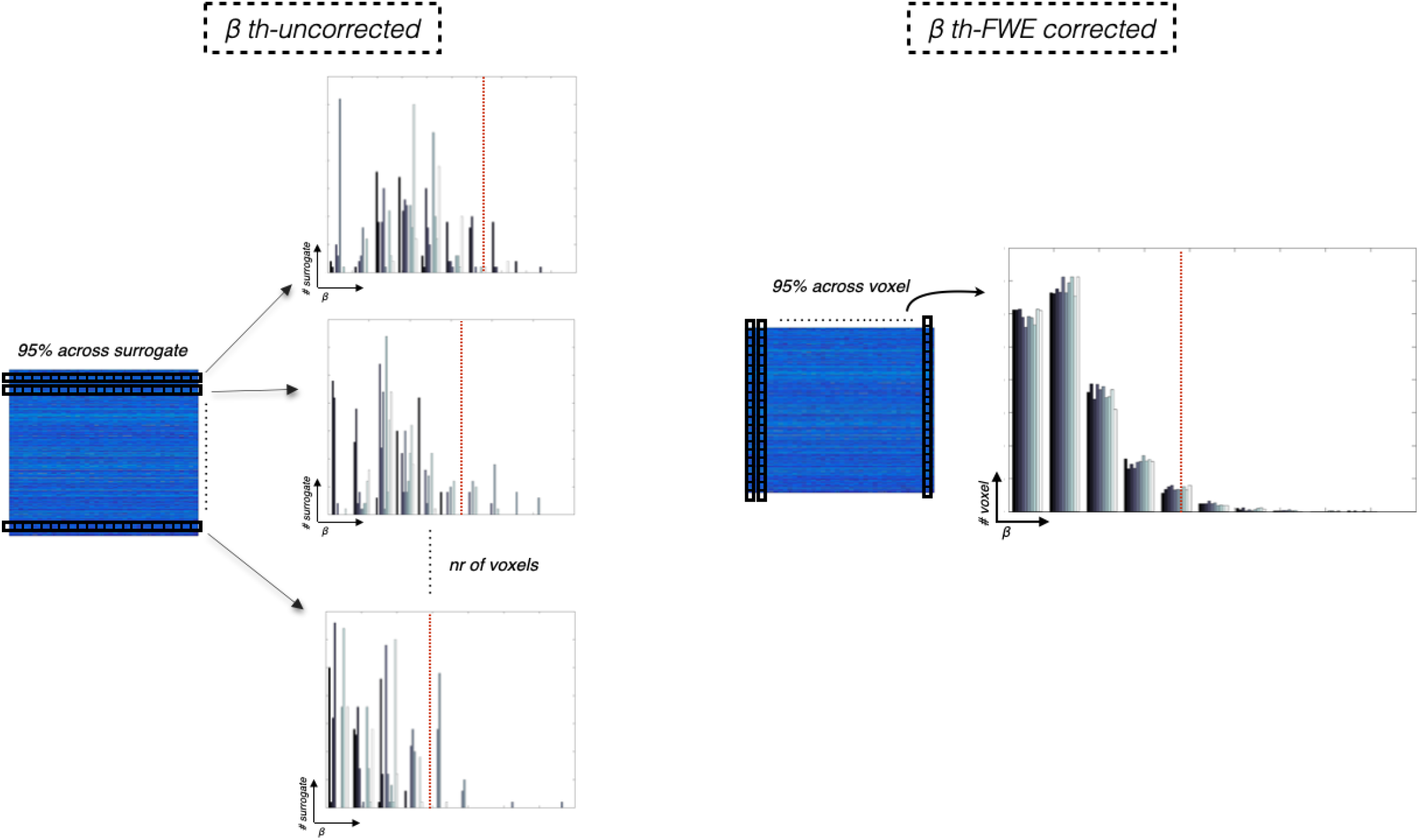
Overview of the *beta* thresholding approach. Using the MCMC CF model (option B), the best CF fit on the surrogate BOLD time series was used to define two different *beta*-thresholds: an uncorrected and a FWE-corrected option. Both *beta*-thresholding options are defined using the 95th percentile estimates of the null distribution, that can be obtained by selecting one pseudo-randomly selected surrogate or averaging all surrogates computed per each voxel. For the first option, an optimal uncorrected threshold is computed for each voxel in the target region (top panel - the optimal uncorrected threshold obtained using the CI option is indicated by red line). Note that the optimal *beta*-threshold may be different for different voxels as it is empirically derived from CF fits to the surrogate BOLD time series of each individual voxel. The second optionis FWE corrected for all voxels in the target regions. In this case, one unique threshold is obtained using the best fit of one pseudo-randomly selected surrogate BOLD time series.

Finally, we compared the voxel selection at the single participant level using both methods and for *beta*-thresholds to the Explained Variance (VE) based on the surrogate data of the MCMC CF model.

#### 2.6.3 Model selection

In order to compare the single gaussian (SG) to the DoG model, two parameters were considered: the classical variance explained (VE) of the model (for illustrative purposes) and the Bayesian Information Criterion (BIC, see equation 17) (Myung & Pitt, 2004; Schwarz, 1978) defined as follows:

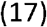

Where is the number of time series and is the number of parameters estimated by the model. Per target voxel, the best model was determined as the one having the lowest BIC value or the highest VE.

## 3. Results

To preview our results, we observe a good level of agreement between the standard CF and the novel Bayesian MCMC CF models. We estimated the uncertainty and (in)dependence for the MCMC CF parameters (and). Moreover, we implemented and showed how to use the new threshold based on the effect size of the model. Finally, we found that a CF model based on a single Gaussian was preferred over one based on a DoG to explain the observed BOLD correlations between visual areas.

### 3.1 MCMC CF model: validation and comparison

Figure 3 compares the standard CF estimates against the MCMC CF (A) estimates by plotting them on a smoothed 3D cortical surface of a representative subject. We directly compared the standard CF and MCMC CF (option A) models as both models estimated the CF parameters using an ordinary least squares fit and retained the same number of parameters. The maps were obtained using V1 as source region while V2, V3, hV4, LO1 and LO2 as target region (Figure 3, panel b and c). Note that the center of a CF is defined at a voxel location in the source region. In order to present results in terms of VFM coordinates, the center position in terms of voxel location is transformed into eccentricity and polar angle values as derived from a pRF mapping. As in the standard CF, the MCMC CF maps show a clear visuotopic organization for all the CF parameters estimated (Figure 3, panel c) and they are in good agreement with the standard CF estimates (Figure 3, panel b and c). The MCMC CF maps obtained for option B are reported in supplementary material (Figure S1).

**Figure 3.**
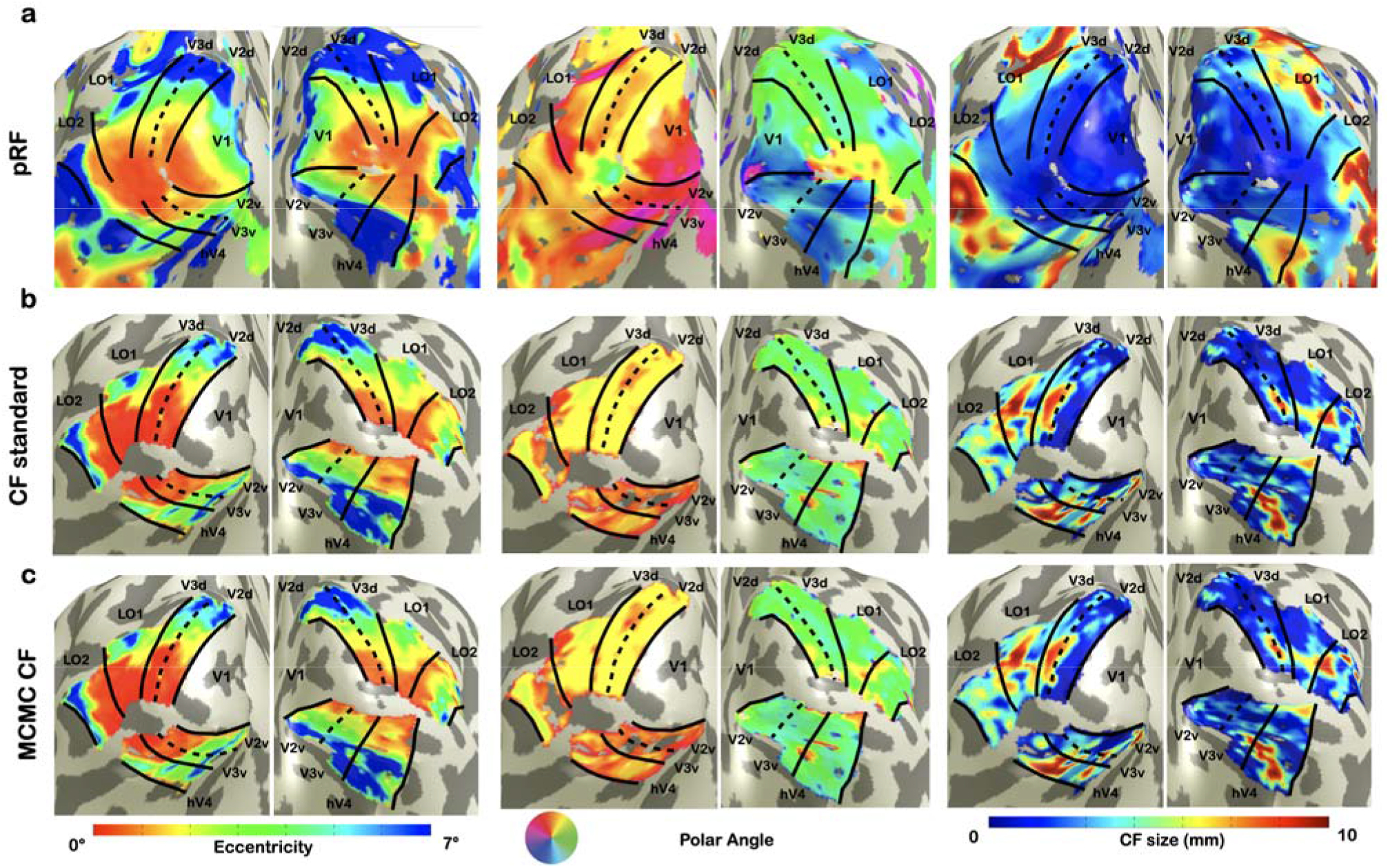
Visualization of pRF and CF maps for a single subject. From left to right: eccentricity, polar angle and pRF/CF size parameters are displayed. Panel **a** corresponds to pRF model estimation and visual area definitions. Panels **b** and **c** show VFM derived maps estimates using CF and MCMC CF (option A) models, respectively.

In line with the earlier work that introduced the standard CF method (Haak et al., 2013), we quantified the differences between the resulting standard CF and MCMC CF maps (in VFM coordinates) by correlating them against the pRF-derived eccentricity and polar angle (Table 2). The same quantification of visuotopic organization at the individual subject level is reported in Supplementary material (Table S1). Overall a good agreement between the two CF approaches was found for the CFs in visual cortex area V2 (i.e. V1 > V2). Given the high degree of correspondence between standard CF and PRF mapping we will use the standard CF maps as a reference from here on.

**Table 2.** Group level correlations between the visual field maps derived using pRF, CF and MCMC CF models. To estimate the level of agreement of CF maps, we computed the correlations coefficients and compared the the eccentricity (*rho*) and polar angle (*theta*) maps obtained using the standard CF and those derived using the MCMC CF models to the pRF *rho* and *theta* (gold standard). P-values based on Spearman’s correlations range from 0.0001 to 0.045 across different rois.

| **ROIs** | **CF** |  | **MCMC CF (A)** |  | **MCMC CF (B)** |  |
| --- | --- | --- | --- | --- | --- | --- |
|  | *rho* | *theta* | *rho* | *theta* | *rho* | *theta* |
| V1 --> V2 | 0.8619 | 0.7328 | 0.8576 | 0.7359 | 0.8570 | 0.7357 |
| V1 --> V3 | 0.8094 | 0.7051 | 0.8075 | 0.7010 | 0.8053 | 0.7068 |
| V1 -> hV4 | 0.7759 | 0.6851 | 0.7698 | 0.6787 | 0.7664 | 0.6966 |
| V1 -> LO1 | 0.7740 | 0.6600 | 0.7627 | 0.6645 | 0.7522 | 0.6677 |
| V1 -> LO2 | 0.5726 | 0.6292 | 0.5800 | 0.6064 | 0.5372 | 0.6212 |

Furthermore, we quantified the level of agreement between the standard CF and the MCMC CF models by correlating the estimated parameters of each model at voxel level per subject (Figure 4). The best level of agreement between all parameters was observed in the early visual areas (Figure 4, panel a, b and c - V1 > V2 and V1 > V3). For some subjects, a weak level of agreement was noticed in V1>LO2 especially if the number of voxels in LO2 is low (Figure 4, V1>LO2). Thus, the good agreement and correspondence observed between the standard CF and the MCMC CF implies that this novel Bayes framework is a valid method to accurately reproduce cortico-cortical properties of the visual field.

**Figure 4.**
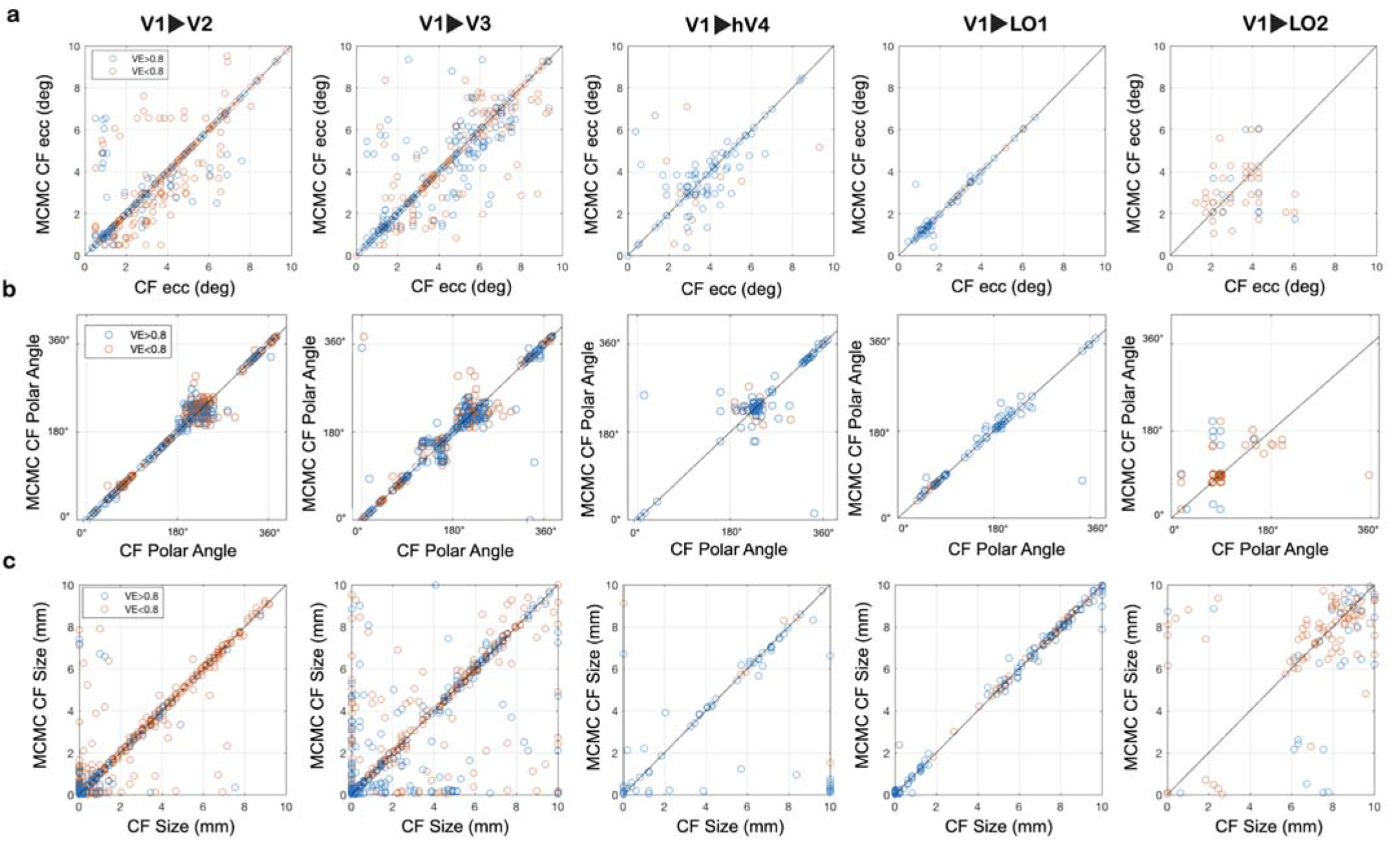
Correlation between CF and MCMC CF parameters at the subject level. From left to right: parameter estimates for CFs in target regions (V2, V3, hV4, L01 and LO2) that sample from V1. Panels **a, b** and **c** show the correlation between CF and MCMC CF models for eccentricity, polar angle (converted into visual field coordinates) and CF size parameters, respectively. Only voxels with explained variance higher than 0.5 in the standard CF model are considered. The remaining voxels were color coded based on the VE. Voxels with VE between 0.5 and 0.8 are color-coded in orange while voxels with VE higher than 0.8 are plotted in blue.

### 3.2 MCMC CF Uncertainty

Based on a quantile difference between _*3*_and _*3*_of the estimated posterior distribution, we estimate a voxel-wise uncertainty (see Equation 16) for the MCMC CF parameters (and). Only the MCMC CF parameters directly estimated using the model (,,VE) are included in the analysis. To evaluate the possible dependency between the MCMC parameter estimates, the corresponding (posterior) uncertainty and the residual noise under a OLS setting, the cross-correlation coefficients between these estimates were computed (Figure 5). Negative cross-correlations were obtained between the model variance explained (VE) and the uncertainty estimate for and irrespective of the target ROI (Figure 5, panel a and b). A weak positive cross-correlation is present between the MCMC CF parameters and the corresponding uncertainties. This indicates that uncertainty is a new, independent parameter. Similar patterns were observed across all ROIs (Figure 5, panel a and b).

**Figure 5.**
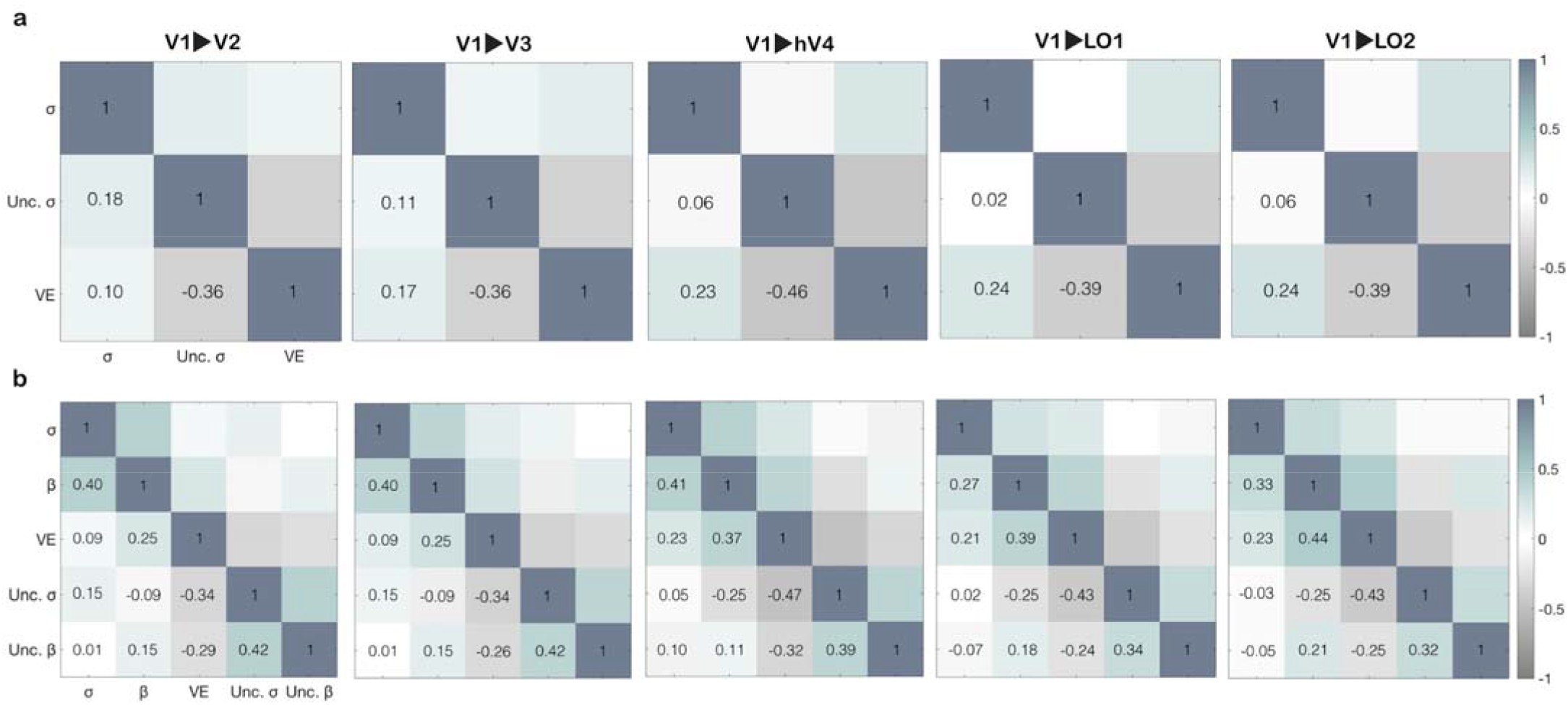
Dependency between MCMC CF parameters and uncertainty at voxel level. Cross-correlations were computed between the estimated MCMC CF parameters and the uncertainty derived from them. Only the CF parameters directly estimated using the model (, and VE) are included in this analysis. Panel **a** shows the dependency obtained from MCMC CF (option A) while panel **b** shows the dependencies for MCMC CF (option B) model.

### 3.3 Beta thresholding

Traditionally, the standard way to threshold voxels is based on the VE which indicates the goodness of fit. Alternatively, a threshold can be obtained from the effect size estimate. Analogous to traditional fMRI analysis, the effect size (*beta*) is compared against its distribution under the null hypothesis of no effect. We implemented two options: 1) an uncorrected threshold and 2) a threshold that is FWE corrected for all voxels in a target ROI.

Figure 6 shows the distributions of value estimated for a single V2 voxel using surrogate (Figure 6, panel b) and real fMRI time-series (Figure 6, panel a). Using the 95th percentile of the null distribution, we defined the *beta*-uncorrected threshold (Figure 6, panel c) and the FWE *beta*-corrected threshold (Figure 6, panel d).

**Figure 6.**
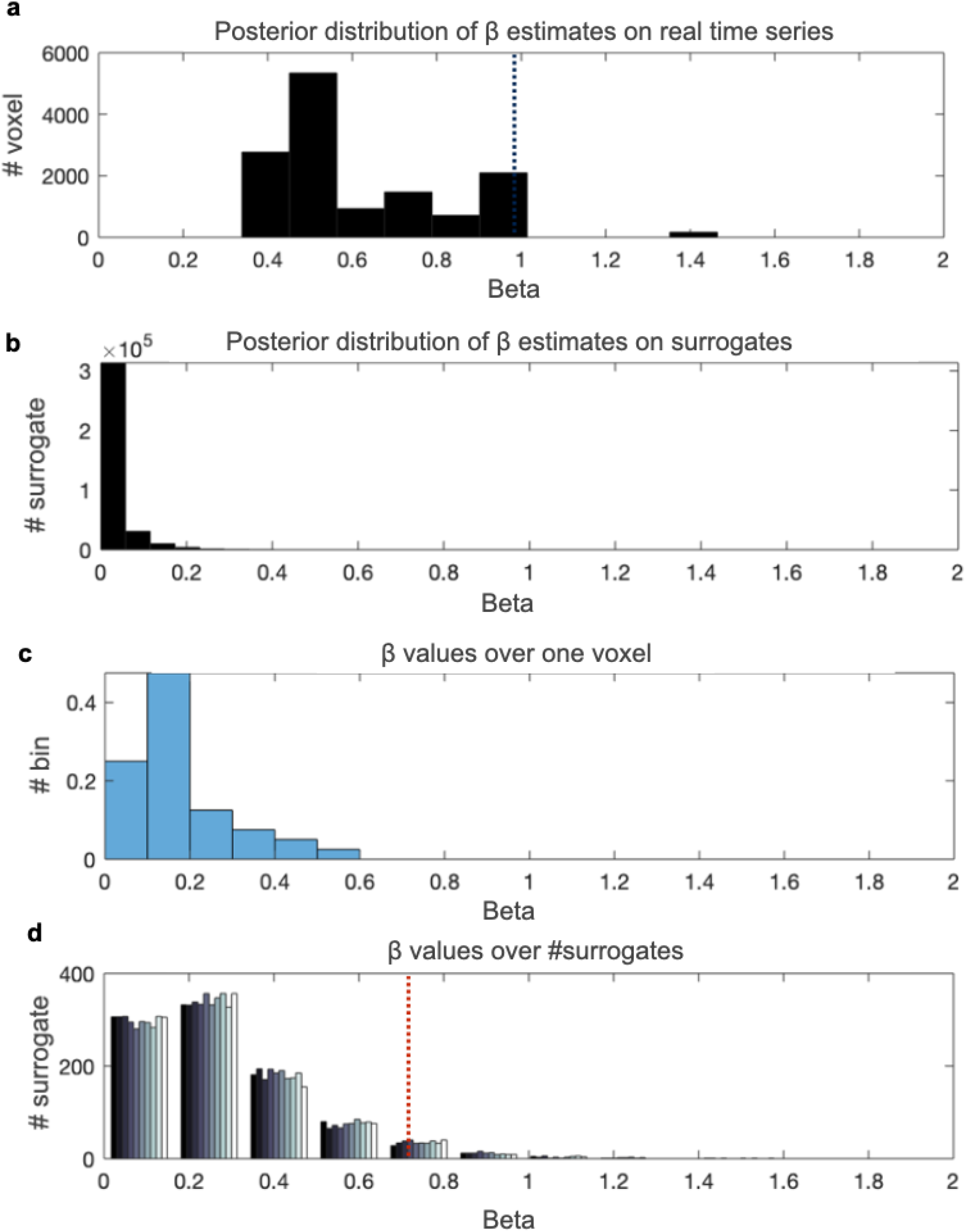
*Beta*-based threshold estimates in a single visual area (V1>V2) for an individual subject. Panels **a,** and **b** show the histogram of posterior distribution obtained from the MCMC procedure for 1 voxel in V2 sampling from V1 using actual and surrogate data, respectively. In panel **a,** the best fitting **MCMC** *beta* estimate for this voxel (= 0.9815) is indicated by a black dotted line. Panel **b** reports the histogram of the posterior distribution under the null hypothesis. This histogram is obtained by combining posterior MCMC data across all 40 surrogates. Panel**c** shows the histogram of the best fitting *beta* values across all 40 surrogates This histogram can be used to obtain an uncorrected threshold. Finally, panel **d** shows the distribution of optimal *beta* values under the null hypothesis (i.e. surrogate data) across all voxelsV2. Each bar represents one surrogate. For visualization purposes, we only present the first ten. The FWE-corrected *beta* threshold (panel **d)** is indicated by a red dotted line.

From here on, the FWE-corrected *beta* threshold obtained using the 95th percentile is applied in the model comparison analysis. A full evaluation of the thresholding method can be found in the supplementary material.

### 3.4 Model comparison

We quantify the goodness of fit for a single Gaussian (SG) and difference of Gaussian (DoG) models by using the standard VE and then, by computing and comparing the BIC scores. For both model predictions, we used MCMC CF option B (see Method section 2.5 and 2.5.1). Only voxels surviving the subject-wise FWE beta-threshold were considered for further analysis.

An increased VE is present for the DoG and this effect is consistent across all visual areas (Figure 8, panel a) albeit that this difference was not significant (1000 permutations were performed; p-values range from 0.6412 to 0.2794 across different rois). An increased VE for the DoG model is expected due to additional degrees of freedom (i.e. more free parameters to be estimated) than the SG model (Haefner & Cumming, 2008; Singh & Horn, 1999). Furthermore, the VE metric is highly biased from the intrinsic variability and residual noise in the data.

**Figure 8.**
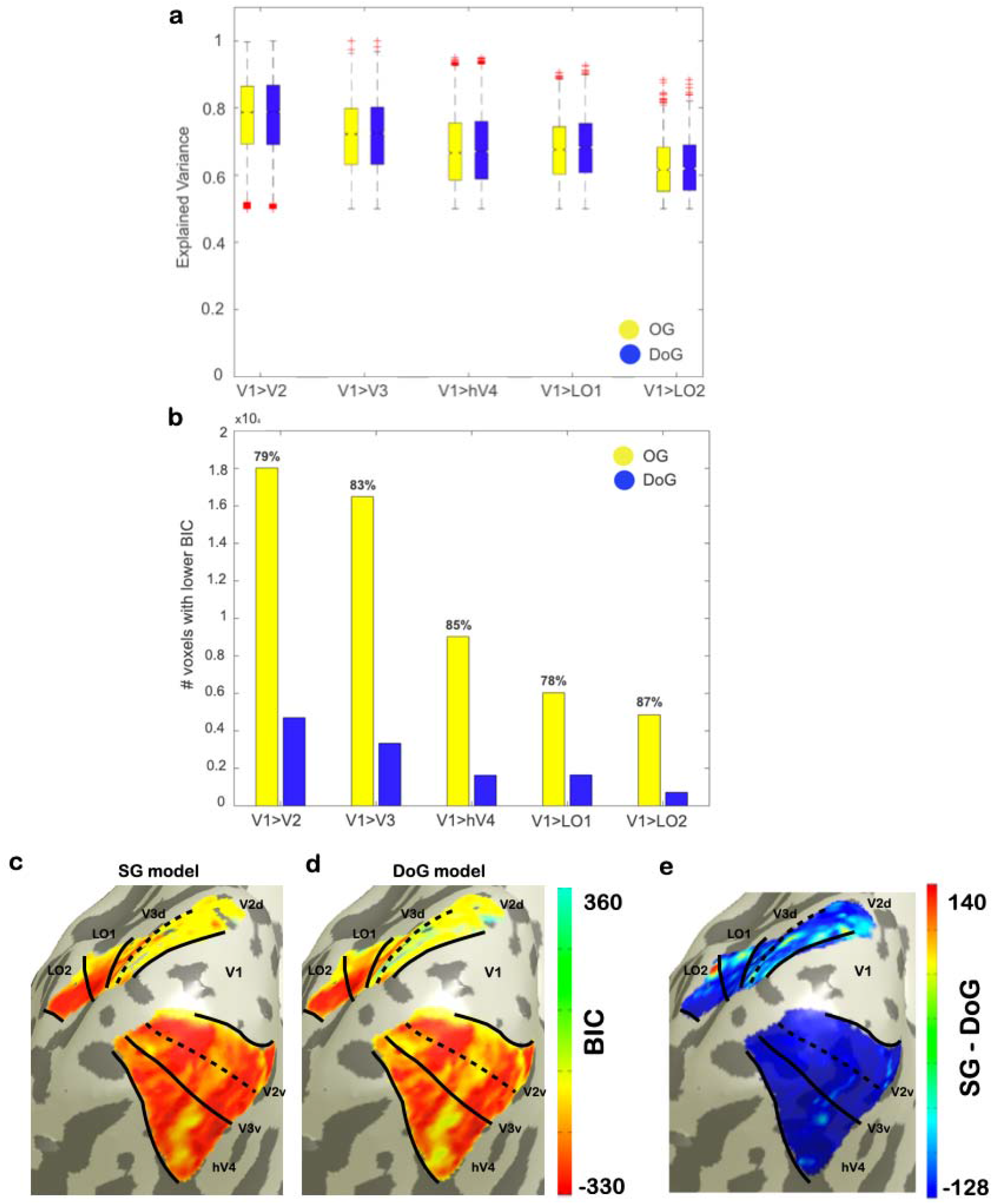
Model comparison between SG and DoG CF models. Variance explained and BIC was computed at single subject level and then grouped over the population per ROI for both SG and DoG MCMC CF models. Data was thresholded using the CI corrected beta-based option. Panel a: the variance explained for both models is reported. In almost all ROIs, the DoG CF model (blue) has an increased VE compared to SG CF model (yellow). While in panelb, the percentage of voxels with lower BIC per single ROI is reported (back). It is notable that the simpler model is favored for all the ROIs in the visual cortex when the number of parameters is also taken into account. Panels **c** and **d** show BIC derived maps estimates using OG and DoG MCMC CF models, respectively. While Panel **d** shows the BIC difference between the two CF models (SG - DoG).

Though the likelihood estimation of the model obtained by using the MCMC CF approach, we computed a Bayesian model estimation (BIC) which enables a proper model comparison between the two models. In contrast, the SG model shows a consistently lower BIC for the majority of voxels (~ 80%) across all visual areas (Figure 8, panel b). Furthermore, we projected the BIC values obtained at voxel level onto a smoothed 3D mesh at single subject level (Figure 8, panel c and d) to investigate possible visuotopic (re-)organization or model preferences per single visual area.

In this case, no visuotopic organization was observed for both SG and DoG MCMC models.

## 4. Discussion

We presented a new Bayesian CF modeling framework based on Markov Chain Monte Carlo sampling. We compared the MCMC CF and the standard CF models and observed a good level of agreement between them. We further showed how the Bayesian CF model provides novel parameters to explore underlying cortico-cortical properties of the visual system. We estimated the uncertainty associated to the estimation of CF size () and effect size (). Subsequently, we implemented and tested a new threshold based on and show how it can be used as a complementary threshold to the VE of the model. Finally, we implemented the spatial center-surround feature of receptive fields at the cortical level and compared its performance to the two-dimensional circular symmetric gaussian variant (SG). Interestingly, at the cortical level, we find that the SG model is prefered over the DoG. Below, we discuss our results in more detail.

### 4.1 The MCMC connective field approach compares well to the standard approach

The novel MCMC CF model provides visuotopic maps qualitatively similar to those obtained using the standard CF model (see Figure 3). Quantifying the similarity between MCMC and classical CF models, we observed the best level of agreement in the early visual areas - V1 projecting to V2 (V1>V2) and V1 projecting to V3 (V1>V3). These quantitative and qualitative results are in agreement with those presented previously (Haak et al., 2013), see Figure 4 and Table 1). This high correspondence validates the Bayesian approach as a useful method to characterise cortico-cortical receptive field properties of visual cortex accurately.

In the past years, different Bayesian approaches have been successfully applied to the pRF method allowing estimation of the the full posterior distribution associated to each of the receptive field properties (Adaszewski et al., 2018; Benson & Winawer, 2018; Carvalho et al., 2020; Quax et al., n.d.; Zeidman et al., 2018). Similar to the Bayesian pRF methods (Quax et al., n.d.; Zeidman et al., 2018), the proposed MCMC CF framework provides the marginal distribution associated with each of the cortico-cortical model parameters. Thus, we can estimate the variability on the data by estimating the uncertainty and possible dependencies between the parameter estimates. By looking at the cross-correlation between the MCMC CF parameters and the corresponding uncertainties, the same relation was observed across all visual cortical areas (Figure 5): at most a weak correlation was found between the MCMC CF parameters and the corresponding uncertainties (correlation < 0.25). Therefore, uncertainties can be treated as additional, independent CF parameters that quantify their reliability. As expected, an inverse relationship was observed between the goodness of fit of the model (VE) and the uncertainty of and. Nonetheless, additional information on the data can be obtained using the Bayes CF framework as will be discussed in the next paragraph.

### 4.2 The effect size provides a sensible threshold

Using our Bayes CF framework, we considered several thresholding techniques based on the posterior distributions of the effect size,. The standard and widely used approach to threshold voxels is based on the explained variance (VE) of the CF model (Haak et al., 2013; Halbertsma et al., 2019), which indicates the goodness of fit of the model. However, high explained variance does not always correspond with low variability in the estimates (Thielen et al., n.d.). In fact, a model fitted to noisy data may still get a high explained variance. Here, we propose an alternative threshold obtained from the effect size estimate compared against its distribution under the null hypothesis. We compared two options versus the standard VE-based approach: 1) an uncorrected threshold and 2) a threshold, FWE corrected for all voxels in a target ROI. Notedly, some voxels with high uncertainty (and therefore high variability) are still selected based on VE thresholding while they are discarded using the other two (uncorrected) thresholds (Figure 7). An FWE-corrected *beta* threshold when obtained using the 95th percentile is the most conservative threshold. In general, both *beta*-thresholds are more sensitive in the selection of voxels compared to the standard VE. Furthermore, our data-driven thresholds are participant-specific. Therefore, these new *beta*-thresholds are expected to be especially useful when the MCMC CF model is applied in clinical populations (e.g with a lesioned visual pathway or brain neurodegeneration). However, these new *beta*-threshold approaches require a proxy distribution for the null hypothesis based on multiple surrogate BOLD time series per voxel. At present, obtaining this is computationally demanding and time consuming. This limitation is much reduced when a FWE corrected threshold is applied in which case a single surrogate BOLD time serie per voxel will suffice. From figure 6, panel d, it can be seen that the distribution of the null hypotheses (across voxels) is consistent for all 40 surrogates tested. Although a FWE corrected threshold is generally considered to be concervative, note that our FWE corrected threshold is still more sensitive than the standard VE and, in contrast to VE, participant-specific.

### 4.3 The spatial center-surround estimation in the visual cortex

Another significant extension of Bayes CF framework is the ability to compare different cortical receptive field models. We compared the single circular Gaussian (SG) model used in the standard CF (Haak et al., 2013), to the Difference of Gaussian (DoG) model that can estimate the center-surround configuration of a population of neurons throughout the visual cortex (Rodieck, 1965; Zuiderbaan et al., 2012). A similar center-surround configuration is widely used in the population RF model (DoG pRF) (Anderson et al., 2017; Grigorescu et al., 2003; Zhang et al., 2009). In pRF modeling, an increased explained variance was reported for the DoG pRF model in the early visual cortical areas (Zeidman et al., 2018; Zuiderbaan et al., 2012). Therefore, they concluded that the DoG pRF model provides a better characterization of the fMRI signal in the visual cortex.

We tested whether a similar argument could be made for CF modeling. We compared the SG and DoG CF models using both the explained variance and the BIC values. Similar to the pRF model comparison, an increased variance explained was reported for the DoG CF model in all the visual cortical areas, albeit the increase was small (see Figure 8, panel a). However, by examining the BIC values, the SG CF model shows a consistent lower BIC for the majority of the voxels (~ 80%) for all visual areas (see Figure 8, panel b). This indicates that the SG CF model is favoured, and that the more complex DoG model may overfit the data. Unlike pRF model, the CF model does not appear to possess a spatial center-surround organisation at cortico-cortical interactions level.

### 4.4 Limitations

Compared to the conventional CF model the Bayes CF framework is computational intense and time consuming. Due to the intense computational load, we decided to use 17500 iterations. This allowed the model to reach the optimal-fit in a reasonable time. Future advances in hardware together with software optimization should contribute to reducing the computation time. Currently, we address this issue by using parallel GPU computing and implementing the method using cluster computation (Avesani et al., 2019).

### 4.5 Future directions

The novel Bayes CF framework presented here uses a straightforward biological-grounded model to assess the cortical receptive field properties, and provides a starting point for future studies.

Similar to the standard CF model, the MCMC CF model is stimuli-agnostic. Previous studies have shown that the standard CF still reflects the visuotopic organization of the visual cortex when applied to BOLD activity recorded in the absence of external stimulation (i.e. resting-state fMRI data) (Bock et al., 2015; Gravel et al., 2014). In a similar way, the MCMC CF model should be able to extract the connectivity based on intrinsic activity. Thus, it can be used to evaluate the quality of cortical processing in participants in which the visual input may be compromised by ocular or neurological lesions.

Besides the estimation of uncertainty, additional benefits can be derived by estimating the full posterior distribution using the MCMC CF method. This will allow to monitor progression of a disease and/or the effect of an intervention by comparing the posterior distribution with the posterior surrogate distribution at a single subject level. This will provide new insights to the underlying cortical mechanisms of neuro-ophthalmic diseases, e.g. glaucoma.

For the first time, the new Bayes CF framework allows to investigate cortico-cortical interactions by using different model definitions. In this study, we tested for the first time the CF DoG model on healthy subjects. This model may provide novel information to characterize inhibitory cortical mechanisms in ophthalmic patients. Besides the DoG model, the MCMC CF can be extended to other model definitions that could possibly characterize additional properties of cortical interaction between visual areas (i.e. elongated shape, rotations, (Zeidman et al., 2018). Furthermore, our framework enables proper cortical model comparison by estimating the likelihood estimation of the model. This allows to compute Bayesian model estimations i.e. BIC, AIC or Bayes Factor at voxel level that can be projected onto a smoothed 3D mesh and further used to investigate possible visuotopic (re-)organization or model preferences per single visual area at single subject level.

## 5. Conclusion

In this study, we have presented and validated a Bayesian inference framework for the CF model using a Markov Chain Monte Carlo approach. We compared the MCMC CF performance to the standard CF model and observed a good agreement using empirical stimulus-driven fMRI data. Novel MCMC CF parameters were included: first, the parameter uncertainty that quantifies the variability and reliability associated with each of the CF parameters. Second, the effect size of the BOLD fluctuation (*beta*) that has been introduced as a reliable, data-driven threshold. Moreover, we implemented different CF kernels. Our results show that a simple SG model is favored in describing the cortico-cortical interactions of the early human visual system. Our new Bayesian CF framework provides an improved and comprehensive tool to assess the neural properties of cortical visual processing that will help to further improve our understanding of the ongoing processes involved in perception, cognition, development, and ageing in both health and disease.

## Acknowledgements

We want to thank Hinke N. Halbertsma for data collection and pRF data analysis.

## Supplementary Material

### Difference of Gaussians: MCMC CF option A

As for MCMC CF model option B described in section 2.5.1, the same MCMC CF fitting procedure described in section 2.5 was used. In the MCMC CF model (option A), both and_*2*_parameters were retained. For each of the two Gaussian distributions, a predicted weighted fMRI signal_*1*_()and _*2*_()was created respectively. In order to orthonormalize both _*1*_()and _*2*_()signals, the Gram-Schmidt process was applied and the inner product between _*1*_()and _*2*_()was consequently used in the OLS fit. For this DoG MCMC CF model, the initial values of _*2*_and_*2*_ were set to 5 and 10, respectively.

### Evaluation of beta thresholding approaches

To evaluate the novel thresholding methods in the voxel selection, we consider the (posterior) uncertainty that is related to residual noise of the model, associated with each MCMC CF parameter, respectively. For each of the threshold techniques, the following thresholds were used: the VE obtained on the null and effect size survived the 95th percentile (for more details, see Method 2.6.2). By comparing both *beta*-uncorrected and *beta*-corrected thresholds (95th percentile) to the standard VE of the null, we noticed that most voxels with high uncertainty are discarded using the beta-threshold approaches and not the null-based VE (see Figure S2, supplementary material). Therefore, the FWE-corrected *beta* threshold obtained using the 95th percentile is applied in the model comparison analysis.

### Supplementary Figures

**Figure S1.**
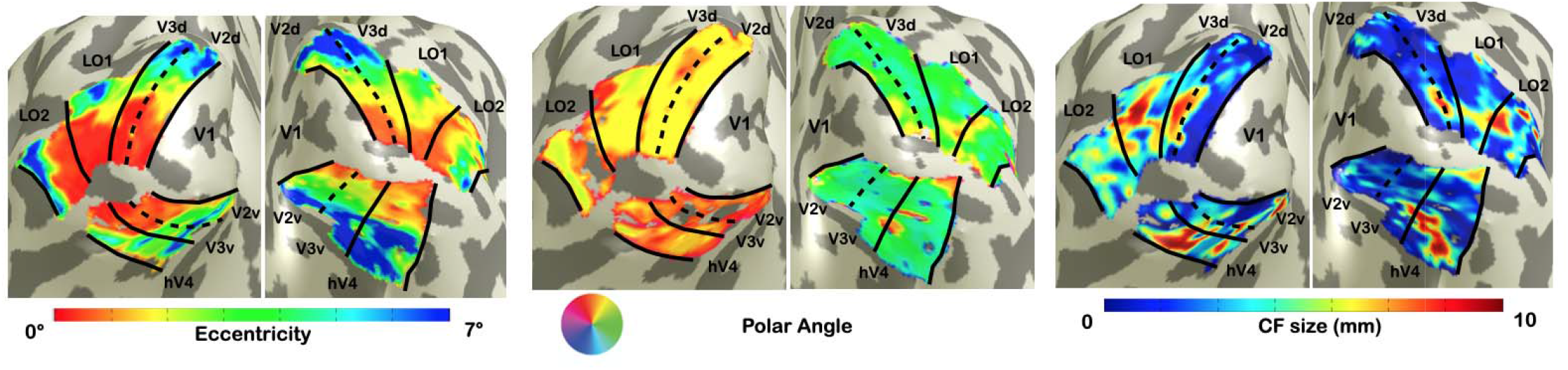
Visualization of MCMC CF (option B) maps for a single subject. From left to right: eccentricity, polar angle and CF size parameters are displayed for MCMC CF (option B) model.

**Figure S2.**
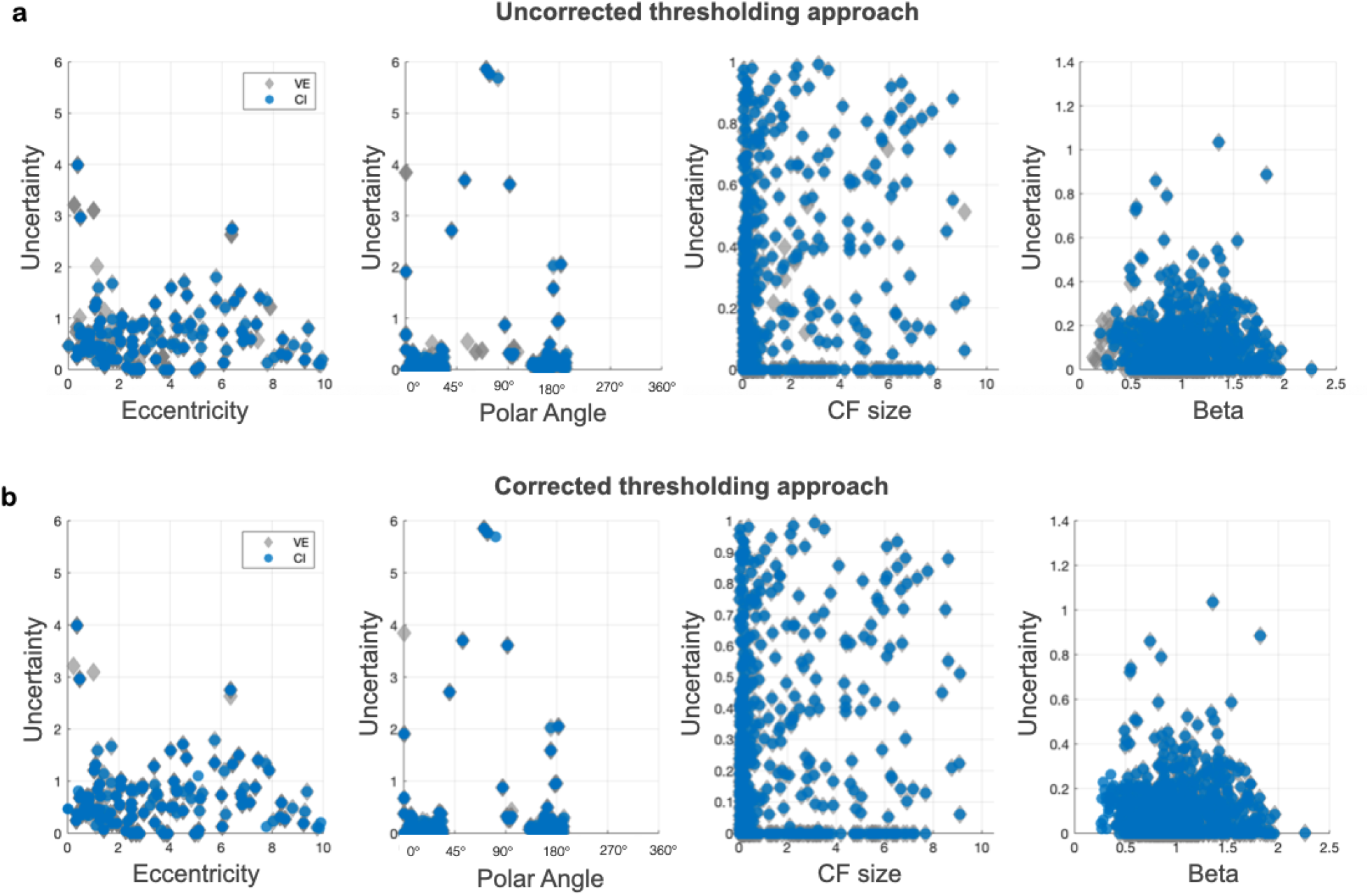
Comparison of thresholding approaches in a single subject (V1>V3). From left to right: estimates of eccentricity, polar angle, CF size and beta parameters and their corresponding uncertainty estimates are displayed, respectively. Only voxels whose beta and VE estimates exceed the 95th percentile of the null were included. In Panel **a,** the *beta*-uncorrected thresholds obtained with the 95th percentile technique were applied and compared to the standard VE of the model. While in Panel **b** the FWE beta-corrected threshold obtained using the 95th percentile technique was applied and compared to the VE.

**Table S1.** Comparison between pRF, CF and MCMC CF parameters at single subject level. To estimate and compare the level of agreement of CF maps, we computed the correlations between eccentricity (*rho*) and polar angle (*theta*) parameters obtained using the standard CF and MCMC CF models to the pRF *rho* and *theta* (gold standard).

Correlations between CF and MCMC CF on VFM data
| V1 > V2 | CF |  | MCMC CF (A) |  | MCMC CF (B) |  |
| --- | --- | --- | --- | --- | --- | --- |
|  | *rho* | *theta* | *rho* | *theta* | *rho* | *theta* |
| sub1 | 0.813 | 0.9124 | 0.8134 | 0.9165 | 0.8065 | 0.9071 |
| sub2 | 0.8407 | 0.9257 | 0.8236 | 0.9269 | 0.8147 | 0.9325 |
| sub3 | 0.9083 | 0.9341 | 0.9073 | 0.9298 | 0.8983 | 0.9212 |
| sub4 | 0.7952 | -0.8866 | 0.7857 | -0.9171 | 0.7814 | -0.9036 |
| sub5 | 0.9256 | 0.9738 | 0.9206 | 0.9738 | 0.9182 | 0.9724 |
| sub6 | 0.912 | 0.8218 | 0.9131 | 0.8726 | 0.9109 | 0.8619 |
| sub7 | 0.8271 | 0.8665 | 0.8244 | 0.8676 | 0.8298 | 0.8465 |
| sub8 | 0.9025 | 0.9231 | 0.9029 | 0.9195 | 0.901 | 0.9301 |
| sub9 | 0.8925 | 0.9247 | 0.8917 | 0.9297 | 0.8744 | 0.9396 |
| sub10 | 0.8606 | 0.4851 | 0.8579 | 0.4887 | 0.8711 | 0.512 |
| sub11 | 0.8757 | 0.937 | 0.8728 | 0.9357 | 0.8812 | 0.9405 |
| sub12 | 0.8171 | 0.9002 | 0.8154 | 0.897 | 0.814 | 0.8979 |
| V1 > V3 | CF |  | MCMC CF (A) |  | MCMC CF (B) |  |
|  | *rho* | *theta* | *rho* | *theta* | *rho* | *theta* |
| sub1 | 0.7624 | 0.8196 | 0.7657 | 0.8254 | 0.7542 | 0.8175 |
| sub2 | 0.6335 | 0.9101 | 0.6339 | 0.9093 | 0.6436 | 0.9261 |
| sub3 | 0.8918 | 0.8146 | 0.8916 | 0.809 | 0.8949 | 0.8434 |
| sub4 | 0.8051 | 0.8123 | 0.7848 | 0.7394 | 0.7708 | 0.649 |
| sub5 | 0.849 | 0.9212 | 0.8437 | 0.9237 | 0.8514 | 0.9318 |
| sub6 | 0.8968 | 0.9579 | 0.8893 | 0.9564 | 0.8925 | 0.9662 |
| sub7 | 0.8039 | -0.7392 | 0.8074 | -0.7291 | 0.8154 | -0.7797 |
| sub8 | 0.8554 | 0.8837 | 0.8572 | 0.8603 | 0.8536 | 0.8908 |
| sub9 | 0.8936 | 0.8486 | 0.8952 | 0.8769 | 0.8754 | 0.8643 |
| sub10 | 0.8219 | 0.6488 | 0.8195 | 0.617 | 0.8142 | 0.5826 |
| sub11 | 0.8289 | 0.6754 | 0.8283 | 0.6675 | 0.829 | 0.778 |
| sub12 | 0.7185 | 0.9349 | 0.7191 | 0.9347 | 0.711 | 0.9276 |
| V1 > hv4 | CF |  | MCMC CF (A) |  | MCMC CF (B) |  |
|  | *rho* | *theta* | *rho* | *theta* | *rho* | *theta* |
| sub1 | 0.7172 | 0.9246 | 0.7195 | 0.9299 | 0.7188 | 0.9418 |
| sub2 | 0.5778 | 0.1933 | 0.5429 | 0.1466 | 0.5467 | 0.1646 |
| sub3 | 0.9082 | 0.4841 | 0.9074 | 0.4747 | 0.8932 | 0.4682 |
| sub4 | 0.8052 | 0.952 | 0.7601 | 0.9549 | 0.704 | 0.9577 |
| sub5 | 0.7429 | 0.5631 | 0.7419 | 0.5511 | 0.7565 | 0.5267 |
| sub6 | 0.8137 | 0.9575 | 0.8221 | 0.9585 | 0.8326 | 0.9302 |
| sub7 | 0.6616 | 0.8195 | 0.68 | 0.8048 | 0.6702 | 0.8506 |
| sub8 | 0.875 | 0.1732 | 0.8765 | 0.1626 | 0.898 | 0.3333 |
| sub9 | 0.8263 | 0.7803 | 0.8272 | 0.787 | 0.8124 | 0.8142 |
| sub10 | 0.8515 | 0.6762 | 0.845 | 0.6709 | 0.8409 | 0.6361 |
| sub11 | 0.8263 | 0.9158 | 0.8286 | 0.9075 | 0.8129 | 0.9127 |
| sub12 | 0.7594 | 0.8641 | 0.7573 | 0.8816 | 0.7645 | 0.8855 |
| V1 > LO1 | CF |  | MCMC CF (A) |  | MCMC CF (B) |  |
|  | *rho* | *theta* | *rho* | *theta* | *rho* | *theta* |
| sub1 | 0.7889 | 0.8991 | 0.791 | 0.8988 | 0.7879 | 0.9116 |
| sub2 | 0.6597 | 0.5441 | 0.5903 | 0.5512 | 0.5486 | 0.5513 |
| sub3 | 0.7639 | 0.9208 | 0.7685 | 0.9013 | 0.7942 | 0.8901 |
| sub4 | 0.6755 | 0.7616 | 0.6754 | 0.7877 | 0.6973 | 0.7399 |
| sub5 | 0.7734 | 0.5203 | 0.7798 | 0.5493 | 0.8 | 0.5553 |
| sub6 | 0.8911 | 0.8911 | 0.8991 | 0.8882 | 0.8954 | 0.8803 |
| sub7 | 0.8128 | 0.9044 | 0.8133 | 0.9067 | 0.7986 | 0.9015 |
| sub8 | 0.8187 | 0.5641 | 0.8251 | 0.5548 | 0.8463 | 0.5802 |
| sub9 | 0.8718 | -0.6411 | 0.8683 | -0.6245 | 0.8524 | -0.6814 |
| sub10 | 0.6833 | 0.7436 | 0.6724 | 0.739 | 0.6213 | 0.7339 |
| sub11 | 0.7746 | 0.8628 | 0.7592 | 0.8636 | 0.7866 | 0.8804 |
| sub12 | 0.7854 | 0.9562 | 0.7823 | 0.9521 | 0.7435 | 0.9223 |
| V1 > LO2 | CF |  | MCMC CF (A) |  | MCMC CF (B) |  |
|  | *rho* | *theta* | *rho* | *theta* | *rho* | *theta* |
| sub1 | 0.3983 | 0.8177 | 0.3658 | 0.8603 | 0.3446 | 0.9007 |
| sub2 | 0.3099 | 0.5619 | 0.3091 | 0.5843 | 0.2575 | 0.6552 |
| sub3 | 0.8312 | 0.7455 | 0.852 | 0.7346 | 0.8186 | 0.8065 |
| sub4 | 0.7063 | -0.4262 | 0.718 | -0.3368 | 0.7169 | -0.5773 |
| sub5 | 0.7415 | 0.8481 | 0.4414 | 0.6072 | 0.419 | 0.6429 |
| sub6 | 0.6265 | -0.2412 | 0.644 | -0.2207 | 0.6303 | -0.2404 |
| sub7 | 0.7037 | 0.9187 | 0.7035 | 0.9744 | 0.6928 | 0.9022 |
| sub8 | 0.8729 | 0.7493 | 0.9135 | 0.6165 | 0.8751 | 0.7138 |
| sub9 | 0.8011 | 0.4694 | 0.8016 | 0.4398 | 0.7457 | 0.6136 |
| sub10 | 0.6504 | 0.8926 | 0.6691 | 0.9241 | 0.5046 | 0.8767 |
| sub11 | 0.8666 | 0.5665 | 0.6419 | 0.7735 | 0.7173 | 0.3133 |
| sub12 | 0.5648 | 0.6997 | 0.636 | 0.7404 | 0.598 | 0.5381 |

